# Cell adhesions link subcellular actomyosin dynamics to tissue scale force production during vertebrate convergent extension

**DOI:** 10.1101/2021.06.21.449290

**Authors:** Robert J. Huebner, Shinuo Weng, Chanjae Lee, Sena Sarıkaya, Ophelia Papoulas, Rachael M. Cox, Edward M. Marcotte, John B. Wallingford

## Abstract

Axis extension is a fundamental biological process that shapes multicellular organisms. The design of an animal’s body plan is encoded in the genome and execution of this program is a multiscale mechanical progression involving the coordinated movement of proteins, cells, and whole tissues. Thus, a key challenge to understanding axis extension is connecting events that occur across these various length scales. Here, we use approaches from proteomics, cell biology, and tissue biomechanics to describe how a poorly characterized cell adhesion effector, the Armadillo Repeat protein deleted in Velo-Cardio-Facial syndrome (Arvcf) catenin, controls vertebrate head-to-tail axis extension. We find that Arvcf catenin is required for axis extension within the intact organism but is not required for extension of isolated tissues. We then show that the organism scale phenotype is caused by a modest defect in force production at the tissue scale that becomes apparent when the tissue is challenged by external resistance. Finally, we show that the tissue scale force defect results from dampening of the pulsatile recruitment of cell adhesion and cytoskeletal proteins to cell membranes. These results not only provide a comprehensive understanding of Arvcf function during an essential biological process, but also provide insight into how a modest cellular scale defect in cell adhesion results in an organism scale failure of development.

## Introduction

Head-to-tail axis extension is an early and essential step of animal development (Keller, 2002), and genes that govern axis extension are directly linked to human birth defects, in particular neural tube closure defects (Wallingford et al., 2013). While the plan for axis extension is encoded in the genome, execution of this program is a mechanical process involving the highly reproducible and tightly coordinated movement of cell collectives (Mongera et al., 2019). Several collective cell movements contribute to axis extension, including convergent extension (CE), in which a group of cells move towards each other along one axis resulting in extension of the perpendicular axis (Keller, 2002; Tada and Heisenberg, 2012). CE is deeply conserved in evolution and has been studied in organisms ranging from nematodes to vertebrates (Huebner and Wallingford, 2018).

Biomechanical analyses have become increasingly critical to developmental biology, especially for understanding morphogenetic processes such as collective cell migration (Barriga et al., 2018), neuronal guidance (Thompson et al., 2019), and organogenesis (Tao et al., 2019). Because axis extension requires tissue scale force to push the animal’s anterior (presumptive head) away from its posterior (presumptive tail), the biomechanics of both elongation generally and CE specifically have been explored in recent years (Mongera et al., 2018; Shook et al., 2018; Xiong et al., 2018),(Keller, 2012). These forces originate from actomyosin contractility within individual cells and the cellular scale forces must be integrated and transduced across the cell collective by cell-cell adhesions (Lecuit et al., 2011). Understanding how cell adhesion links cellular scale force production to tissue scale biomechanics, especially in vertebrates, is an ongoing challenge.

Classical cadherins are the key adhesion molecules that connect cell-cell contacts to the actomyosin cytoskeleton (Charras and Yap, 2018). Extensive biochemical, structural, and cell biological studies have characterized the proteins connecting cadherin-based adhesions to cellular force production machinery, but these data primarily apply to cultured cells or epithelial tissues (Charras and Yap, 2018). Critically, however, cadherin dependent CE in early vertebrate embryos also occurs in mesenchymal tissues and much less is known about cadherin function during collective movement of mesenchymal cells (Theveneau and Mayor, 2012; Walck-Shannon and Hardin, 2014). Thus, to understand early vertebrate development it is essential to improve our comprehension of cadherin-based cell adhesion in mesenchymal tissues.

Cadherins have been shown to control biomechanics across multiple length scales during CE (Huebner et al., 2020; Kale et al., 2018), but cadherin function absolutely requires an array of effector proteins, and less is known about the effector proteins controlling biomechanics in mesenchymal tissues. One poorly characterized cadherin effector, the catenin Armadillo Repeat protein deleted in Velo-Cardio-Facial syndrome (Arvcf) is required for multiple developmental processes, including head-to-tail axis extension (Cho et al., 2011; Fang et al., 2004), but the biomechanical underpinnings of this defect are unclear. At the molecular level, Arvcf is most closely related to p120-catenin and similarly is thought to control cadherin trafficking (McCrea and Park, 2007). Arvcf is also associated with Rho and Rac activity and thus has a role in signaling to the actomyosin cytoskeleton (Fang et al., 2004).

Here, using a combination of classic embryology, biomechanical measurements, and cell biology, we found that Arvcf specifically modulates the force generated by CE but interestingly not the ability of a tissue to converge and extend, per se. Rather, this embryonic phenotype results from a failure of convergent extension to generate sufficient force to push an animal’s anterior away from its posterior. We further show that this force generation defect is associated with a modest reduction in cell adhesion and a more pronounced dampening of the oscillatory recruitment of cadherin and actin to cell membranes. These results not only provide a deeper understanding of a poorly defined catenin during an essential biological process, but also illustrates that a modest change in cell behavior can result in an organism scale failure of development.

## Results

### Identification of tissue and stage specific Cdh3 protein interactions in *Xenopus* convergent extension

The classical cadherin Cdh3 (aka C-cadherin; aka mammalian P-cadherin) plays a crucial role in morphogenesis of the dorsal marginal zone (DMZ) mesoderm of the *Xenopus* gastrula (Fagotto et al., 2013; Huebner et al., 2021; Lee and Gumbiner, 1995; Pfister et al., 2016), a powerful and deeply studied paradigm for understanding CE (Keller and Sutherland, 2020). Cadherins control numerous cellular processes including proliferation, cell migration, cell polarity, and mechanotransduction and to achieve this assortment of cellular tasks, cadherins interact with an array of structural and signaling molecules (Arslan et al., 2021). However, we currently lack a comprehensive roster of proteins interacting with Cdh3 in the *Xenopus* DMZ. We therefore used a combination of classic embryology and modern mass spectrometry to determine the protein interactome for Cdh3.

Details of our affinity purification mass spectrometry (AP-MS) approach can be found in the Methods Section, but briefly, we used mRNA injection to express Cdh3-GFP or GFP alone in the *Xenopus* dorsal mesoderm and then isolated DMZ (Keller) explants at the onset of gastrulation (~st.10.25)(Keller, 2012)(Fig. 1A). Explants were allowed to undergo CE *ex vivo* until mid-gastrulation (~st.12) and were lysed. Cdh3-GFP or GFP alone were purified by GFP pulldown and associated proteins were identified by mass spectrometry (Fig. 1A), using the GFP-only pulldown as a negative control. This experiment was performed in two biological replicates with ~800 explants per condition, per replicate.

**Figure 1:**
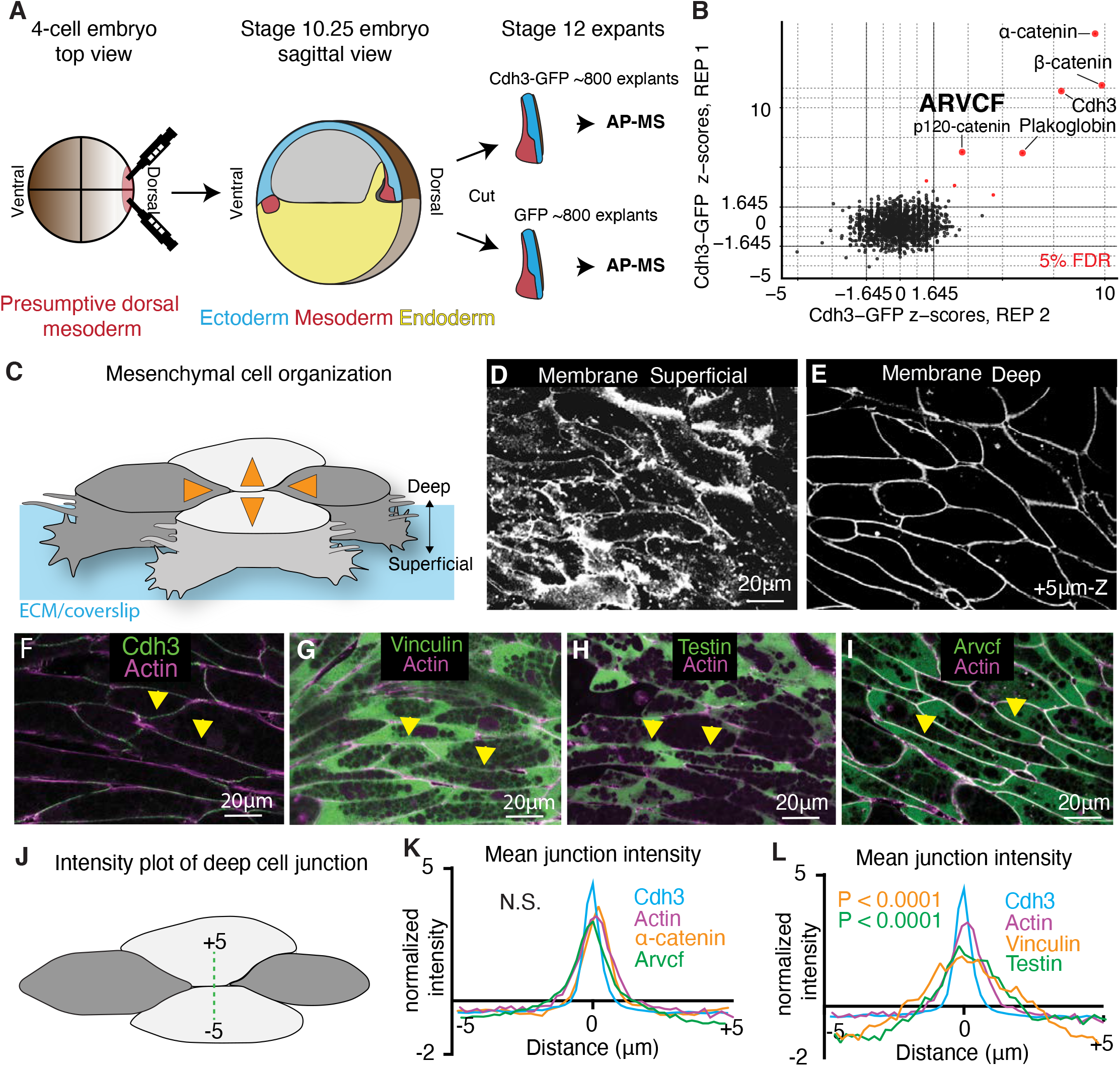
Tissue and stage-specific Cdh3 affinity purification mass spectrometry results in a highly specific Cdh3 protein interaction dataset. **A.** Schematic depicting the method used for tissue and stage-specific affinity purification mass spectrometry (APMS) of Cdh3. Targeted microinjection was used to introduce Cdh3-GFP or GFP alone, a negative control, into the presumptive dorsal mesoderm of 4-cell stage *Xenopus* embryos. Microdissection was used to excise the dorsal mesoderm and the dorsal ectoderm during early gastrulation (~ St.10.25) (Keller explants). Explants then converged and extended until mid-gastrulation (~ St.12) at which time explants were lysed and GFP-pulldown was used to purify for Cdh3-GFP or GFP alone. Purified proteins were prepared and sent for mass spectrometry. **B.** Graph showing the relative protein orthogroup enrichment (see methods) from two replicates of the Cdh3 AP-MS experiment. Here we plot the z-scores from replicate 1 on the y-axis and the z-scores from replicate 2 on the x-axis. Z-scores are a statistical comparison of proteins detected in the Cdh3-GFP pulldown versus the negative control divided by the standard error from the null distribution based on total peptide counts. Each dot represents a protein identified in the Cdh3 AP-MS dataset and red dots represent proteins that fall below a 5% FDR threshold. **C.** Cartoon depicting *Xenopus* mesenchymal cells during convergent extension. Here the mediolateral (ML) cells, dark gray, move to each other resulting in displacement of the anterior-posterior (A-P) cells, light gray. Orange arrows show the cell movements. The mesenchymal cells become elongated along the ML axis at the onset of CE and acquire polarized mediolateral oriented cellular protrusions. Mesenchymal cells show apparent structural differences along the superficial (cell-ECM interface) to deep (cell-cell interface) axis. Here polarized lamellar-like structures are observed at the superficial surface. Movement deeper into the cell reveals cell-cell interfaces and actin-based protrusions which extend between neighboring cells. **D.** Image of the superficial surface of converging and extending *Xenopus* mesenchymal cells. Cells are labeled with a membrane marker which primarily shows lamellar-like protrusions at the cell-ECM interface. **E.** Image of the deep surface of the same cells shown in Fig. 1D. Here the membrane marker largely highlights the cell-cell junctions. **F.** Image of Cdh3 (green) and actin (magenta) at deep cell-cell contacts. Yellow arrowheads point to cell junctions. **G.** Image of vinculin (green) and actin (magenta) at deep cell structures. Yellow arrowheads point to cell junctions. **H.** Image of testin (green) and actin (magenta) at deep cell-cell junctions. Yellow arrowheads point to cell junctions. **I.** Image of the Arvcf (green) and actin (magenta) at deep cell-cell junctions. Yellow arrowheads point to cell junctions. **J.** Schematic displaying the method used to measure fluorescent intensities at cell-cell interfaces. **K.** Intensity plots of Cdh3 (blue), actin (magenta) α-catenin (orange), and Arvcf (green). Here zero is set at the center of the cell-cell junction and each protein shows a clear peak at the cell junction. Each line is the average over dozens of line plots from a minimum of three replicates. Distributions were statistically compared using a KS test. **L.** Intensity plots of Cdh3 (blue), actin (magenta), vinculin (orange), and testin (green) at cell-cell junctions. Vinculin and testin lack peaks at the cell-cell junction and the distributions of vinculin and testin were statistically different from Cdh3 and actin as compared by a KS test. Each line represents the average of dozens of line plots over a minimum of three replicates.

Because of the genome duplication in the *X. laevis* genome (Session et al., 2016), we identified enriched proteins by collapsing homeologs and highly related entries into eggNOG vertebrate-level orthologous groups, as outlined in the Methods. Z-scores and confidence intervals for each protein orthogroup were then calculated based on the ratio of peptide counts in the experimental (Cdh3-GFP) compared to negative control (GFP alone) and the standard error from the null distribution based on the global peptide counts. (For a complete list of proteins identified, peptide counts, Log2 fold change over GFP alone, z-scores, p-values, and confidence intervals, see Data Table 1.)

Crucially, our analysis identified Cdh3 itself and the well-known cadherin interactors α-catenin and β-catenin as the most highly enriched proteins in our dataset (Fig. 1B). Using the same parameters (joint z-score over both replicates with a multiple hypothesis-corrected p-value <= 0.05), we also identified less well known but important Cdh3 interactors, including Plakoglobin (aka JUP) (Aktary et al., 2017) and the orthogroup containing the related catenin proteins Arvcf and p120 (McCrea and Park, 2007) (Fig. 1B)(Supp. Fig.1). Taken together, these results demonstrate that our AP-MS dataset effectively identified Cdh3 interacting proteins in the *Xenopus* gastrula DMZ.

### Localization of Cdh3-ineraction partners reveals similarities but also differences from the E-cadherin/Cdh1 paradigm

Cells engaged in CE in the *Xenopus* gastrula DMZ display a unique organization, and previous work has identified two sites of Cdh3 action. Unlike more familiar epithelial cells, which have clear apico-basal polarity, these mesenchymal cells lack a well-defined molecular asymmetry along the axis spanning the superficial cell-extracellular matrix (ECM) interface and the deeper cell-cell junctions (Green and Davidson, 2007)(Fig. 1C). The superficial surface is visually dominated by large actin based lamellar-like structures that become polarized to the mediolateral cell edges during CE (Fig. 1C, D), and Cdh3 localizes to foci in these protrusions (Pfister et al., 2016) (Supp. Fig. 2A). Observation deeper into the tissue (3-5μm) reveals clear cell-cell junctions (Fig. 1C, E), though extensive actin-based protrusions between neighboring cells at these deep cell-cell interfaces are also observed (Green and Davidson, 2007; Weng et al., 2021); Cdh3 localizes to dynamic foci at deeper cell-cell interfaces (Fagotto et al., 2013; Huebner et al., 2021). Because of the key role of these deeper cell-cell interfaces in CE (Huebner et al., 2021; Shindo et al., 2019; Shindo and Wallingford, 2014; Weng et al., 2021), we used live imaging to identify Cdh3-interacting proteins that localize to these sites.

As mentioned, Cdh3 localizes strongly to the deep cell-cell junctions, where it co-localized with actin, as expected (Fig. 1F; quantified in Fig. 1K-L). Given our interest in biomechanics, we next examined the localization of the mechanosensitive protein α-catenin, a very strong Cdh3 interactor in our AP-MS data (Fig. 1B). α-catenin displayed a similar localization at deep cell-cell junctions (Fig. 1K). This result is consistent with the requirement of α-catenin in zebrafish CE (Han et al., 2016).

A surprising result was that the mechanosensitive protein vinculin was only very weakly enriched in our Cdh3 AP-MS dataset from the *Xenopus* DMZ (Data Table 1). Vinculin is a key component of Cdh1/E-cadherin cell-cell junctions and is required for CE of *Drosophila* epithelial cells (Huveneers et al., 2012; Kale et al., 2018), so we expected to find vinculin at the deep mesenchymal cell-cell junctions. Nonetheless, we found that vinculin-GFP was not co-localized with Cdh3 and α-catenin at deep cell-cell junctions (Fig. 1G, L). Rather, vinculin-GFP localized specifically to the foci in superficially positioned lamellipodia (Supp. Fig. 2B), where Cdh3 is also known to act (Pfister et al., 2016).

This result motivated us to examine another mechanosensitive protein, Testin, which is required for *Xenopus* CE and is a known tension-dependent effector of cadherin adhesion (Dingwell and Smith, 2006; Oldenburg et al., 2015). Like Vinculin, Testin was present but not highly enriched in our AP-MS dataset (Data Table 1), and moreover, Testin-GFP did not localize to deep cell-cell junctions and instead was enriched only in foci in the superficial lamella (Fig. 1H, L; Supp. Fig. 2C).

Finally, amongst the most highly enriched orthogroups in our Cdh3 AP-MS dataset was that containing the closely related Arvcf and p120 catenins (Data table 1)(Fig. 1B)(Supp. Fig. 1). By examining individual peptides in this orthogroup, we found that while both proteins interact with Cdh3, Arvcf had both the larger peptide count and fold change (Supp. Fig. 1). Arvcf is required for *Xenopus* CE (Fang et al., 2004; Paulson et al., 2000), yet little is known of its function, and even its localization during CE has not been reported. Using a functional Arvcf-GFP fusion, we found that Arvcf was present in the superficial lamellar structures (Supp. Fig. 2D, white arrows) but moreover was prominently enriched in the deep cell-cell junctions, where it co-localized strongly with Cdh3 (Fig. 1I, yellow arrows, Fig. 1K).

Together, these localization patterns indicate that the Cdh3-dependent mesenchymal cell-cell junctions in *Xenopus* DMZ cells share some features of the more-thoroughly characterized E-cadherin-dependent adhesions in epithelial cells. However, the absence of vinculin and testin from these junctions suggests important differences as well. These data motivated us to explore cell adhesion during *Xenopus* CE in more detail, and we chose to focus on the Arvcf catenin.

### Arvcf is required for head-to-tail axis extension in the embryo but is dispensable for CE in isolated tissues

The very high confidence interaction of Arvcf with Cdh3 in the *Xenopus* DMZ (Fig. 1B), its requirement for CE, and its connection to Rho and Rac signaling (Fang et al., 2004; Reintsch et al., 2008) made it a strong candidate for further investigation. We therefore used a previously characterized morpholino oligonucleotide to disrupt Arvcf function (Fang et al., 2004). We recapitulated the described axis elongation defect after Arvcf knockdown, and moreover, this defect was rescued with Arvcf-GFP (Fig. 2A-D). As an additional control, we used targeted injections to generate mosaics and showed with immunostaining that Arvcf KD significantly depleted the endogenous Arvcf protein (Fig. 2E-H). Together these data demonstrate the efficacy and specificity of our approach for Arvcf loss of function.

**Figure 2:**
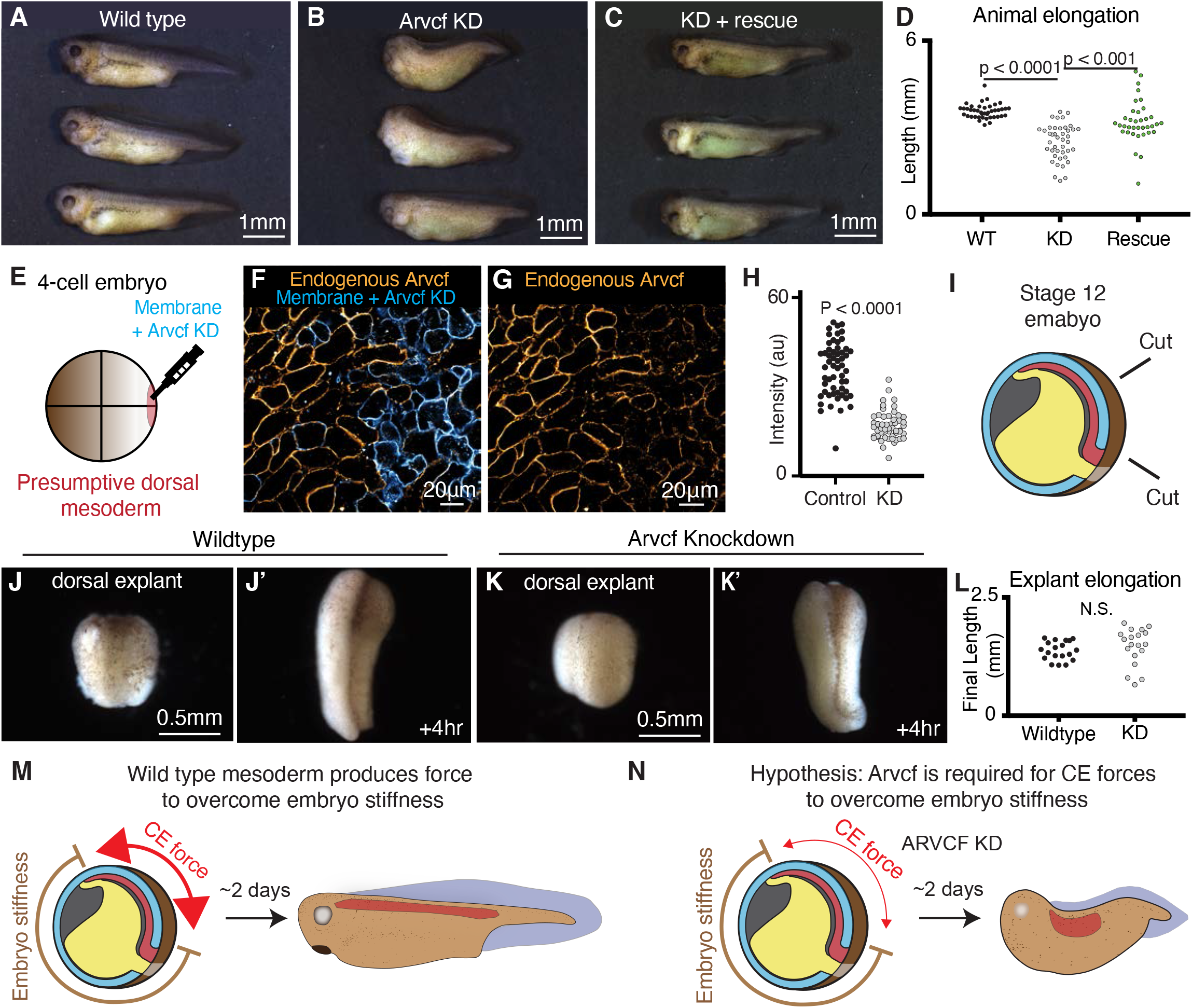
Arvcf is required for embryonic axis extension but is dispensable for CE in explants. **A.** Wild type tadpoles (~st.40). **B.** Sibling embryos to those shown in Fig. 2A in which Arvcf was knocked down in the dorsal mesodermal cells. The Arvcf depleted embryos have a shortened head-to-tail axis and display a characteristic dorsal bend suggesting a CE defect. **C.** Sibling embryos in which Arvcf was knocked down and then rescued with exogenous expression of Arvcf-GFP. **D.** Plot showing tadpole (~st.40) head-to-tail length for the wildtype, ARVDF knockdown, and Arvcf rescue conditions. Embryo lengths were statistically compared using an ANOVA test. Each dot represents a single embryo and data was collected from a minimum of three experiments. **E.** Cartoon depicting the microinjection method used to generate mosaic animals. Here we inject the Arvcf morpholino, along with a membrane marker, into a single dorsal blastomere at the 4-cell stage. This results in mosaic knockdown of Arvcf in approximately half of the dorsal mesodermal cells. **F.** Immuno-staining for Arvcf (orange) in an embryo in which Arvcf has been mosaically knocked down (blue cells). **G.** The same image shown in Fig. 2F except the membrane marker has been removed to better visualize the Arvcf immunostaining. Arvcf protein levels are reduced in the cells that received the morpholino. **H.** Quantification of endogenous Arvcf protein levels from the immunostaining performed on embryos with mosaic Arvcf knockdown. The wildtype cells are shown in black and knockdown cells are shown in gray. Each dot represents the average ARVCF intensity at the membrane of a single cell and data was collected from a minimum of three replicates. Conditions were statistically compared using a Students T-test. **I.** Cartoon depicting the microdissection used to isolate dorsal explants from late gastrula embryos (~st.12) (dorsal isolate). **J.** Image of a wildtype dorsal isolate. **J’.** Image of a wildtype dorsal isolate after a four-hour period of extension. **K.** Image of a dorsal isolate in which Arvcf has been knocked down in the mesodermal cells. **K’.** Image of an Arvcf depleted dorsal isolate after a four-hour period of extension. Here the Arvcf depleted explant elongated to approximately the same extent as wild type explants. **L.** Graph showing the final dorsal isolate length, after a four-hour elongation, for wildtype and Arvcf depleted embryos. Each dot represents the length of a single explant and conditions were statistically compared using a Student T-test. **M.** Cartoon depicting the forces involved in *Xenopus* axis extension. Here the dorsal mesoderm (red) and the overlaying neural ectoderm (blue, above red) converge and extend generating force to push against the stiff embryo. In the case of wildtype embryos, the CE generated force is sufficiently large (red arrows) to overcome the embryo stiffness and the resulting animals have elongated head-to-tail axis. **N.** We hypothesize that Arvcf is required for CE generated force and that depletion of Arvcf reduces the tissue level force produced by CE. In this case the reduced CE force is insufficient to push the stiff surrounding embryo and axis extension fails. One prediction of this hypothesis is that Arvcf deficient explants would not show an elongation defect when there is no external resistance, which fits with our data reported in Fig. 2J-L.

The failure of axis extension, and in particular the dorsal bend, observed in the Arvcf knockdown embryos (Fig. 2B) strongly suggested a CE defect. As a more direct test of the role of Arvcf in CE, we used a traditional explant assay in which converging and extending tissue is excised from the embryo and allowed to extend *ex vivo* (Fig. 2I). For this assay, the entire dorsal region of the embryo was excised at late gastrula stage (~st.12) (dorsal isolate) and allowed to elongate for 4 hours to early neurulation (~st.14). To our surprise, we observed no difference in the extent of elongation when comparing the Arvcf knockdown dorsal explants with control explants (Fig. 2J-L).

To understand the embryonic phenotype of Arvcf loss of function, we considered that axis elongation in *Xenopus* embryos requires increasing force generation of the converging and extending dorsal tissues to overcome the stiffness of surrounding embryo (Zhou et al., 2009; Zhou et al., 2015). We therefore hypothesized that Arvcf was not in fact required for CE *per se*, but instead was required for sufficient CE-mediated force generation to overcome resistance from the surrounding embryo (Fig. 2M, N). To test this idea, dorsal explants were isolated at late gastrula stages (~st. 12), embedded in a semi-compliant 0.3% agarose gel and incubated for 4 hours (Fig. 3A). In this constrained condition, control explants elongated effectively (Fig. 3B, B’, D), but under the same conditions, we observed a significant reduction in the extension of explants deficient for Arvcf (Fig. 3C-D). Thus, by simply adding a subtle mechanical challenge to the Arvcf deficient explants, we were able to recapitulate the embryonic Arvcf knockdown phenotype. This result suggests that Arvcf controls force production during vertebrate CE.

**Figure 3:**
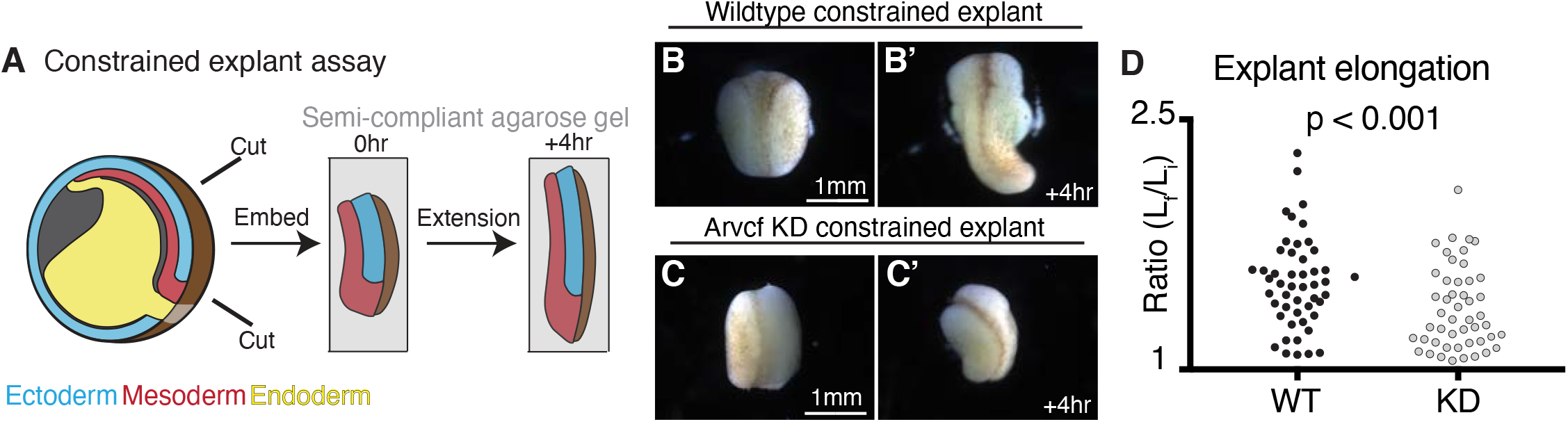
External constraint of Arvcf deficient explants recapitulates the embryonic axis extension defect. **A.** Schematic depicting the constrained explant assay used to mimic the mechanical environment experienced within the embryo. Here dorsal explants were excised from late gastrula (~st.12) embryos (dorsal isolate). Explants were then embedded in a simi-compliant 0.3% low melting temperature agarose gel that approximates the stiffness of the embryo. Images were acquired after embedding the explants and then a second round of images was acquired after the explants elongated over a 4-hour interval. **B.** Image of a dorsal isolate after embedding in gel. **B’**. Image of the same dorsal isolate shown in Fig. 3B after 4 hours of elongation. **C.** Image of an Arvcf deficient dorsal isolate after embedding. **C’**. Image of the same dorsal isolate in Fig. 3C after a 4-hour interval during which the explant had minimal elongation. **D.** Graph showing the extent of dorsal isolate elongation during the constrained explant assay for wildtype and Arvcf knockdown dorsal isolates. The y-axis shows the ratio of the final explant length over the initial explant length. Each dot represents a single dorsal isolate and conditions were statistically compared using a Students T-test.

### Arvcf is required for tissue-scale force production and stiffening during convergent extension

Our data suggest a specific role for Arvcf in the generation of strong mechanical forces by CE, so we next used a biomechanical assay to measure the force output of wildtype and Arvcf deficient dorsal isolates. Following the methods developed by (Zhou et al., 2015), we embedded explants dissected from late gastrula embryos (~ st.12) in semi-rigid, 0.6% agarose gels containing evenly dispersed fluorescent beads (Fig. 4A). Within this semi-rigid gel, wildtype explants converged and produced an extending force that deformed the gel (Fig. 4B, B’). We also observed tissue buckling at ~3 hours when a sudden bend appeared (Supp. Fig. 3A, B, arrow). The embedded fluorescent beads provided fiducial markers for tracking gel deformation, allowing us to observed compression in the gels at the AP poles of the explant (Fig. 4B’). In contrast, Arvcf KD explants also converged and buckled (Fig. 4C), but gel deformation by CE was nearly unobservable (Fig. 4C’).

**Figure 4:**
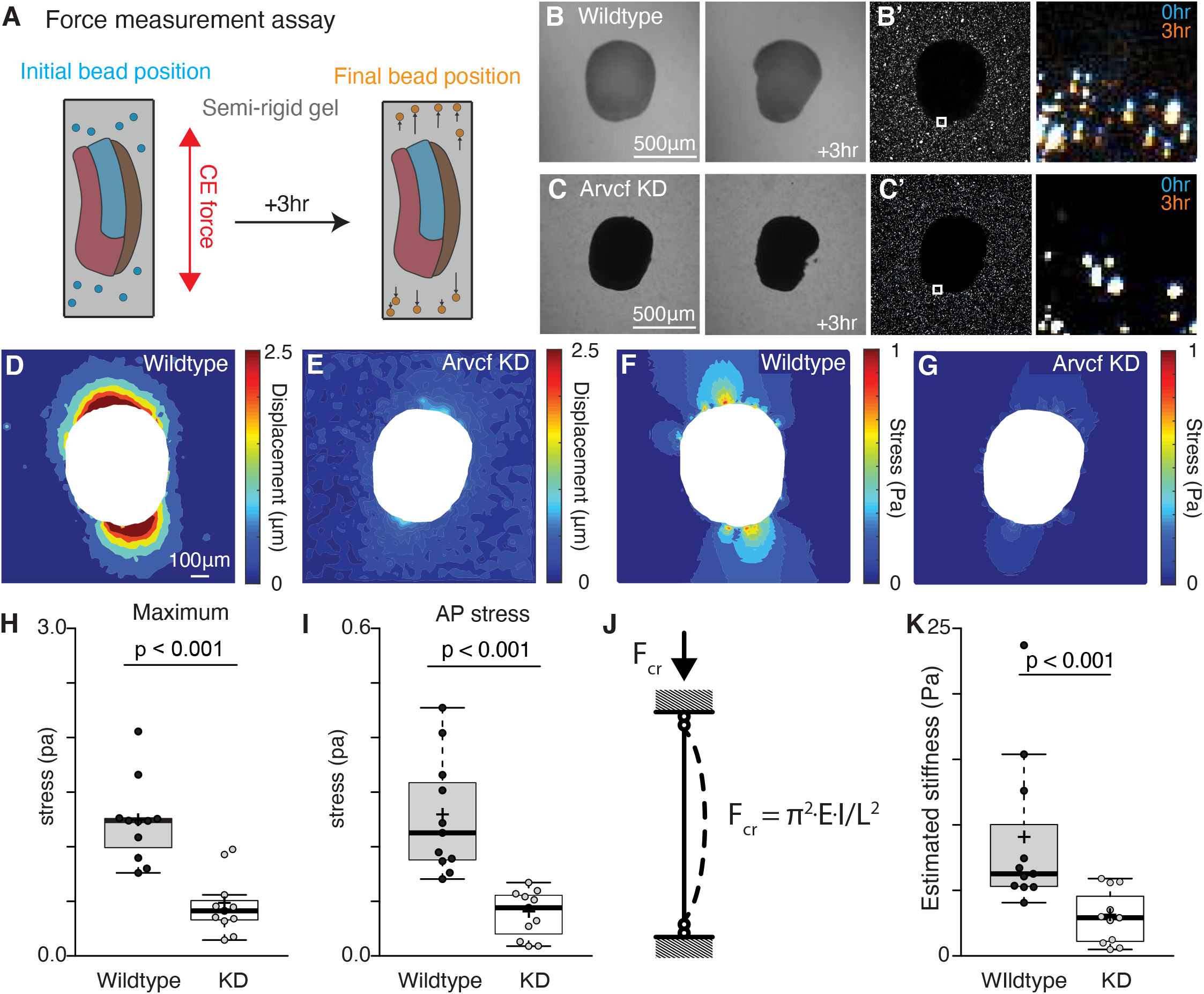
Arvcf controls force production during vertebrate convergent extension. **A.** Schematic depicting the assay used to measure the CE force production. Dorsal explants were excised from late gastrula embryos (~st.12) and embedded in semi-rigid 0.6% agarose gels with known mechanical properties. Fluorescent beads were also embedded in gel to allow visualization of the gel deformation. Explants were then incubated for three hours to allow CE. Then with the known mechanical properties of the gel and the displacement field of the beads we calculated the stress fields generated by each explant. **B.** Images of a wildtype explant embedded in a semi-rigid agarose gel and then a second image of the same explant after a three-hour incubation. White arrow points to the direction of the out-of-plane explant buckling. **B**’. Image of the same explant and gel shown in Fig. 4B but here we are visualizing the beads embedded in the gel and surrounding the explant. The inset focuses on the beads adjacent to the explant and the zoomed in image shows both the initial bead position (blue) and the final bead position (orange). **C.** Image of an Arvcf depleted explant embedded in a semi-rigid agarose gel and a second image of the same explant after a three-hour incubation. **C’**. Image of the beads surrounding the explant shown in Fig. 4C. The inset focuses on a subset of beads next to the explant and the zoomed image shows the initial bead position (blue) and the final bead position (orange). **D.** Displacement field measured by PIV from the beads in Fig. 4B’. **E.** Displacement field measured by PIV from the beads in Fig. 4C’. **F.** Von Mises stress field estimated using finite element method in the gel shown in Fig. 4B. **G.** Von Mises stress field estimated using finite element method in the gel shown in Fig. 4C. **H.** Graph showing the maximum compressive stress along the explant-gel interface. Conditions were statistically compared using a Student T-test and Arvcf depleted explants applied a significantly lowered force on the gel. **I.** Graph showing the average compressive stress along the AP axis. Conditions were statistically compared using a Student T-test and Arvcf depleted explants applied a significantly lowered extending force along the AP axis. **J.** Schematic depicting the buckling model to estimate tissue stiffness. Explant was modeled as a column with a rectangular cross-section. When it converged and extended in a semi-rigid gel, the reactive force applied a uniform longitudinal load that caused an out-of-plane tissue buckling, **K.** Graph showing the estimated tissue stiffness using a simplified buckling model. Conditions were statistically compared using a Student T-test and Arvcf depleted explants were significantly softer.

To quantify the compressive force generated by elongating explants, we used particle image velocimetry (PIV) to quantify the movement of fluorescent beads, and thus gel deformation (Fig. 4D, E). Using finite element analysis to estimate the stress field (Fig. 4F, G), we compared the maximum tissue force before the explants buckled (~3hr). Wildtype explants generated maximum compressive stress of 1 Pa at the AP poles and average stress of 0.2 Pa along the AP axis (Fig. 4F, H, I). These values are comparable with prior assessment of CE forces in *Xenopus* (Zhou et al., 2015). By contrast, explants deficient for Arvcf generated significantly less force, displaying a roughly 60% reduction in both maximum and mean stress (Fig. 4G, H, I). Finally, we used the critical buckling load to assess the tissue stiffness (Fig. 4J). We performed a rough estimation by treating an explant as a column with an rectangular cross-section, on which tissue scale extending force applied a uniform longitudinal load. This estimation also revealed a significant reduction in the tissue stiffness of explants deficient for Arvcf (Fig. 4K). These data demonstrate that Arvcf is not only required for tissue scale force production but also for tissue stiffening during CE.

### Arvcf knockdown disrupts cell adhesion, but not cell polarization during convergent extension

To better understand the nature of the Arvcf phenotype, we next examined the series of well-defined cellular behaviors that drive *Xenopus* mesodermal CE. For example, at early gastrulation cells in the DMZ become robustly polarized, aligning and elongating in the mediolateral axis (Shih and Keller, 1992). However, we observed no difference in cell polarization, as Arvcf-KD cells aligned and elongated normally (Fig. 5A-D).

**Figure 5:**
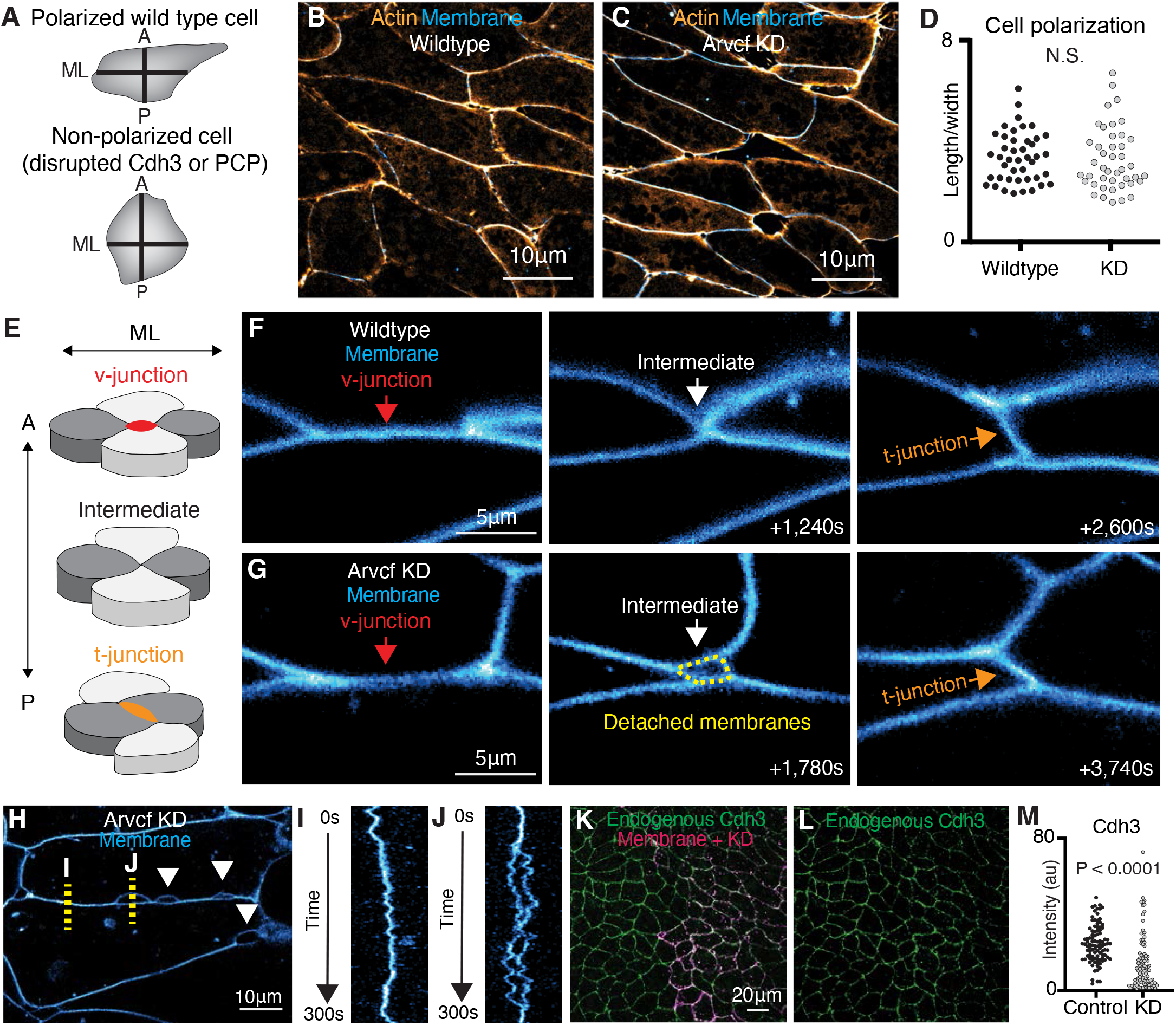
Arvcf is not required for cell polarization but ARVCF knockdown reduces cell-cell adhesion. **A.** Cartoon showing mesodermal cell polarization. These cells elongate along the mediolateral axis during CE and most perturbations of CE, such as depletion of Cdh3 or disruption of PCP signaling, block cellular polarization. **B.** Image of wildtype mesodermal cells displaying characteristic mediolateral cell polarization. Actin is labeled in orange and membrane in blue. **C.** Image of Arvcf depleted cells which also show this characteristic mediolateral polarization. Actin is labeled orange and membranes are blue. **D.** Graph of cell polarization for wildtype and Arvcf knockdown cells. Here polarization was assigned as the ratio of the mediolateral length over the anterior-posterior width. Each dot represents a single cell and there was no statistical difference between the conditions via Students t-test. **E.** Cartoon depiction of the cell movements that drive CE with emphasis on the cell-cell junctions. Initially there is a cell-cell junction between the anterior-posterior cells (light gray) termed a v-junction (red). The cells then intercalate bringing the mediolateral cells (dark gray together. The mediolateral cells then form a new cell-cell contact (t-junction; orange) pushing the anterior posterior cells apart. **F.** Still frames from a time-lapse movie of wildtype cells intercalating. Cell membranes are labeled blue, and the cell intercalation can be visualized as the v-junction is replaced by a t-junction. **G.** Frames from a time-lapse movie showing one example of Arvcf depleted cells intercalating. Here we initially observe a v-junction which shortens, forms a 4-cell intermediate, which then resolves to form a new t-junction. One feature that was unique to the intercalation of the Arvcf depleted cells was that there were often gaps (yellow dashed lines) between the membranes at the intermediate state. Despite these gaps, cells were able to intercalate after ARVCF-KD. **H.** Still frame from a time-lapse movie of cells depleted for Arvcf and expressing a membrane marker (blue). Here gaps are observed between neighboring cells after Arvcf KD, white arrows, suggesting reduced adhesion between cells. **I.** Kymograph showing a 300s time-interval of cell-cell junction dynamics from the yellow dashed line shown in Fig. 5H, labeled I. This kymograph shows a region of membrane without gaps. **J.** Kymograph showing a 300s time-interval of membrane dynamics where a membrane gap opens and then closes. Such membrane gaps are only observed in Arvcf knockout cells. This kymograph is from the time-lapse movie in Fig. 5H, yellow dashed line labeled J. **K.** Immunostaining for endogenous Cdh3 (green) in a field of cells where Arvcf had been mosaically knocked down. A membrane marker (magenta) was used as a tracer for the Arvcf morpholino. **L.** Image showing the isolated Cdh3 channel from Fig. 5K. **M.** Graph displaying the measurement of endogenous Cdh3 intensity from wildtype or Arvcf depleted cells. Each dot represents the average cdh3 intensity of a single cell and conditions were statistically compared using a Students t-test.

Ultimately, tissue elongation is driven by intercalation cell movements that can be simplified to the rearrangement of groups of four cells. Here, anterior-posterior cells are initially connected via a cell junction, commonly called a v-junction, and intercalation of mediolateral cells along the ML axis displaces the v-junction, resulting in formation of a new cell contact, t-junction (Fig. 5E, F). In time-lapse movies, Arvcf-depleted cells were motile and underwent cell intercalation events (Fig. 5G). However, these intercalation events were subtly defective, as the cell membranes often became detached, leaving extra-cellular spaces, particularly as the v-junctions shortened and were replaced by t-junctions (Fig. 5G). Thus, while the junction detachment suggests loose cell attachment in the absence of Arvcf, the end results were generally successful cell intercalation events. A subtle defect in adhesion was also apparent in the altered morphology of cell-cell junctions. For example, we frequently observed gaps in the cell-cell contacts between neighboring cells (Fig. 5H, white arrows) that were not observed at wild type cell-cell junctions. Further, these membrane gaps were highly dynamic with some regions of a membrane remaining connected (Fig. 5I) while gaps opened and closed in other regions (Fig. 5J).

Interestingly the closest homolog of Arvcf, p120-catenin, is known to control cadherin turnover in epithelial tissues (Davis et al., 2003) and loss of Arvcf has a subtle effect on bulk levels of Cdh3 in *Xenopus* (Fang et al., 2004). We therefore, used targeted microinjection to knock down Arvcf in one half of the dorsal mesoderm and used immuno-staining to measure endogenous Cdh3 protein levels (Fig. 5K, L). We measured Cdh3 intensity at cell-cell junctions for wild type and Arvcf depleted cells and found a modest but significant reduction in endogenous Cdh3 protein levels in the knockdown cells (Fig. 5K-M). Together, these results suggest that Arvcf depletion elicits it effects on force generation during CE at least in part by decreasing, but not eliminating, Cdh3-mediated cell adhesion.

### Arvcf depletion does not alter polarized enrichment of actomyosin at v-junctions

Having determined that Arvcf controls tissue-scale force production and cell adhesion, we next asked whether Arvcf regulates polarization of the actomyosin machinery. During *Xenopus* CE, contractile actomyosin is enriched at v-junctions (Shindo and Wallingford, 2014), so we used biosensors for actin (lifeact-RFP) and contractile myosin (myosin light chain 9, Myl9-GFP) to ask if actomyosin localization was altered after Arvcf knockdown. Strikingly, contractile actomyosin appeared to be appropriately localized to the v-junctions in control and Arvcf deficient cells (Fig. 6A, B), and using intensity plots to measure actin and myosin at v-junctions, we observed no difference between controls and Arvcf KD cells (Fig. 6C, D). These data suggest that Arvcf is not required for the steady state polarization of the contractile actomyosin machinery at v-junctions.

**Figure 6:**
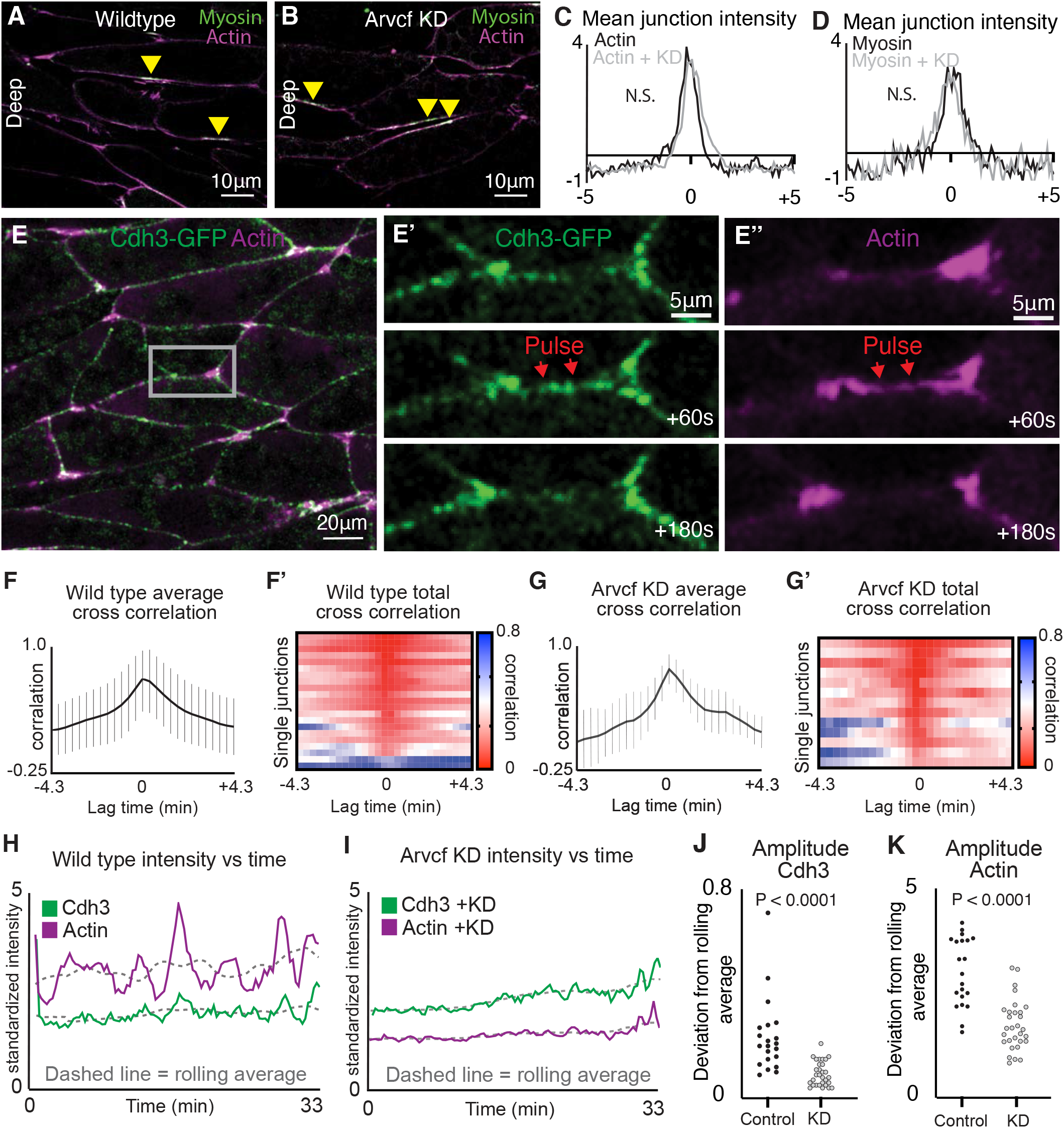
Depletion of Arvcf reduced cell-cell adhesion and disrupted oscillatory temporal dynamics of Cdh3 and actin. **A.** Image of deep cell-cell interfaces showing actin in magenta and myosin in green. Yellow arrowheads point to myosin accumulations at anterior-posterior cell junctions. **B.** Image of deep cell-cell interfaces after Arvcf knockdown. Yellow arrows point to myosin enrichments at anterior-posterior cell junctions. **C.** Intensity plots showing the mean intensity of actin at deep anterior-posterior cell-cell interfaces. Wildtype is shown in black and Arvcf knockdown cells are shown in gray. Conditions were statistically compared using a KS test. **D.** Intensity plots showing mean intensity of myosin at deep anterior-posterior cell-cell junctions. black shows wildtype and gray shows Arvcf depleted cells. Conditions were statistically compared using a KS test. **E.** Frame from a time-lapse movie of wildtype cells, expressing Cdh3-GFP (green) and an actin marker (magenta), during CE. Inset highlights single shortening v-junction. **E’**. Frames from the time-lapse movie shown in Fig. 6E with a zoom on Cdh3-GFP. Here we observe pulsatile enrichment (red arrows) followed by reduction of Cdh3 at shortening v-junctions. **E’’**. The same frames shown in Fig. 6E’ but here showing the actin channel. Actin underwent pulsatile enrichment concurrently with the Cdh3 enrichment. **F.** Graph showing the mean cross correlation between Cdh3 and actin at wildtype shortening v-junctions. Error bars show the standard deviation at each lag time. **F’**. Heatmap displaying the cross correlations for each individual wildtype shortening v-junction. **G.** Graph showing the mean cross correlation between Cdh3 and actin at shortening v-junctions after Arvcf KD. Error bars show the standard deviation. **G’**. Heatmap of the cross correlations of individual shortening v-junctions after Arvcf KD. **H.** Cdh3 (green) and actin (magenta) intensity plotted over time from a wildtype shortening v-junction. Dashed lines show the rolling average (over 3.33 min) for either Cdh3 or actin. **I.** Plot showing Cdh3 (green) and actin (magenta) intensity over time from an Arvcf depleted shortening v-junction. Dashed lines show the rolling average. **J.** We quantified the oscillation amplitude as the deviation of a protein’s intensity plot from that same protein’s rolling average. This graph shows the average Cdh3 amplitude for wildtype and Arvcf knockdown v-junctions. Each dot is the average amplitude from a single shortening v-junction. Conditions were compared using a Students t-test. **K.** The same quantification shown in Fig. 6J but looking at actin oscillation amplitude. Again, each dot represents the average amplitude at a single shortening v-junction and conditions were compared using a Students t-test.

### Arvcf controls the amplitude of pulsatile oscillations of Cdh3 and actomyosin at v-junctions

Cell intercalation in *Xenopus* involves not only polarized steady state enrichment of actomyosin at v-junctions (Shindo and Wallingford, 2014), but also oscillations of actomyosin that correlate with intercalation speed and efficiency (Shindo et al., 2019). Similarly, we recently showed that Cdh3 undergoes oscillatory pulses of clustering that tune the local mechanics of cell-cell junctions (Huebner et al., 2021). We therefore explored the impact of Arvcf loss on the dynamics of actomyosin and Cdh3 clustering during CE.

We collected time-lapse movies of v-junctions during intercalation labeled with Cdh3-GFP and an actin marker (lifeact-RFP). Visual inspection of these movies showed the expected pulsatile enrichment of both Cdh3 and actin, and moreover, these pulses appeared to be coordinated during junction shortening (Fig. 6E, E’’). Moreover, measurement of the cross-correlation of Cdh3 and actin intensity across the junction over time revealed that these proteins underwent coordinated oscillations at shortening v-junctions (Fig. 6F, F’). Importantly, these coordinated oscillations were specific to shortening v-junctions, as such correlation was significantly reduced at non-shortening junctions (Supp. Fig.4). Next, we collected time-lapse images of Cdh3 and actin in Arvcf depleted explants and measured the intensity of these proteins at shortening v-junctions. Cross-correlation analysis showed that Cdh3 and actin pulses remained coordinated at shortening v-junctions in the Arvcf depleted explants (Fig. 6G, G’). Together, these data show that Cdh3 and actin undergo coordinated pulsatile behavior at shortening v-junctions and that Arvcf depletion does not disrupt this coordination.

Strikingly, however, plotting Cdh3 or actin intensity over time provided another means to visualize the pulsatile behavior for wildtype and Arvcf-KD junctions (Fig. 6H, I). Visual inspection of these plots showed a dramatic reduction in the amplitude of the pulses when comparing Arvcf depleted junctions to wildtype (Fig. 6H, I). To compare amplitudes from multiple cell-cell junctions we measured the rolling average intensity for each protein (Fig. 6H, I; dashed lines) and then defined the oscillation amplitude for that junction as the mean deviation between the junction intensity and the rolling average. Using this method, we found a significant difference in the Cdh3 and actin oscillation amplitude when comparing Arvcf-KD cells to wild type (Fig. 6J, K). Thus, Arvcf specifically disrupts the oscillation amplitude -but not coordination- of Cdh3 and actin pulses at shortening v-junctions. In summary, Arvcf-KD results in weaker actin oscillations at v-junctions and we suggest that this reduction in force at the level of individual junctions results in the tissue-scale force defect that causes a failure of embryonic head-to-tail axis extension.

## Discussion

Our over-arching goal here was to explore the role of cadherin-based cell adhesions specifically in the collective movement of mesenchymal cells in a vertebrate embryo. Using proteomic, biomechanical, and cell biological approaches, we identify Arvcf catenin as a key regulator of the biomechanics of CE during *Xenopus* gastrulation. Specifically, we showed that Arvcf was not required for the cell movements that drive CE per se, but instead controlled the force generated by these cell movements. We then show that this phenotype is associated with a modest reduction in cell adhesion and a more substantial defect in the amplitude of the oscillatory recruitment of Cdh3 and actin to cell-cell junctions. We conclude that the low oscillatory recruitment of this actomyosin machinery reduces force production at the cellular level which results in the observed tissue-scale force defect. We feel that several aspects of this study are of note.

First, our Cdh3 AP-MS dataset led us to re-examine the Arvcf catenin, which has known roles in vertebrate development (Cho et al., 2011; Fang et al., 2004) but its cell biological mechanisms of action remain unclear. Our finding that Arvcf specifically controls the magnitude of tissue-scale CE force generation is interesting, as it demonstrates that the force of CE can be uncoupled from dynamic CE cell behaviors. Exactly how Arvcf acts remains unclear, but our work here provides two clues. First, we observed a modest reduction in cell adhesion, likely stemming from a modest loss of junctional Cdh3 (Fig.5), consistent with previous work in *Xenopus* (Fang et al., 2004). The far more significant cell biological phenotype was the reduction of the amplitude of actin oscillations and the amplitude of associated oscillations of Cdh3 (Fig. 6). This result is particularly intriguing because Arvcf (and other catenins in the p120 family) are thought to function quite differently from other catenins. For example, the α-catenin, β-catenin and vinculin complex binds the carboxy-terminus of cadherins and forms a direct linkage to the actin cytoskeleton (Tian et al., 2011). By contrast, Arvcf binds the juxtamembrane region of cadherins to act at least in part via regulation of signaling from the cadherin adhesion complex (McCrea and Park, 2007).

In this light, data from our companion paper provides key insights (see (Weng et al., 2021). CE in epithelial cells in both mice and *Drosophila* is driven by the combined action of basally positioned cell crawling via mediolaterally-positioned protrusions and contraction of apically positioned anteroposterior cell-cell junctions (Huebner and Wallingford, 2018; Sun et al., 2017; Williams et al., 2014). We have now shown that these two modes of intercalation are not physically separated and work in tandem in mesenchymal cells of the *Xenopus* gastrula (Weng et al., 2021). We further show that the integration of the two modes is crucial for high efficiency of cell intercalation. Strikingly, mosaic labelling experiments in that paper suggest that the dampened amplitude of actin oscillations following Arvcf-Kd shown here (Fig. 6) result largely from a specific effect of Arvcf on the mediolateral protrusions (Weng et al., 2021). This result is interesting: On the one hand, it is consistent with the previous finding that Arvcf is required for normal Rac activation during *Xenopus* CE (Fang et al., 2004), and Rac is generally considered to be essential for lamellipodial protrusions (Nobes and Hall, 1995). On the other hand, other work has suggested that disruption of Rac is not sufficient to disrupt lamellipodia formation in *Xenopus* (Tahinci and Symes, 2003). We feel that further work in this area will be exciting.

Regardless of the precise nature of the cell biological mechanisms, our finding that a slight reduction in cellular force was sufficient to disrupt axis elongation in the mechanically challenging environment of the embryo -but not in an isolated, un-constrained tissue- is also important. This result provides new insights into the molecular control of the tissue stiffening that has been well-characterized not only in amphibian embryos (Keller and Sutherland, 2020; Shook et al., 2018; Zhou et al., 2009), but in other vertebrates as well (Mongera et al., 2019). Perhaps most significantly in this light, our data suggest that Arvcf-mediated control of actomyosin and cell adhesion is a central feature of embryos’ “mechanical accommodation” abilities that allow for consistently normal develop despite the wide range of mechanical variation between individual embryos (Zhou et al., 2015).

Finally, in addition to shedding new light on CE, an essential morphogenetic cell movement, our results also provide more general insights into molecular and biomechanical aspects of cadherin-based cell adhesion, specifically in mesenchymal cells. Unlike the more well-characterized cadherin-based junctions in epithelial cells (Rodriguez-Boulan and Macara, 2014), mesenchymal cell junctions are amorphous, lacking the more ordered features such as adherens junctions and tight junctions (Ewald et al., 2012). Nonetheless, mesenchymal cell collectives maintain cohesion during collective movement (Theveneau and Mayor, 2012), including during CE (Walck-Shannon and Hardin, 2014). Thus, we find it striking that vinculin, a key component of Cdh1 (E-cadherin) junctions in epithelia (le Duc et al., 2010), was not only not strongly enriched in our Cdh3 AP-MS dataset, but moreover was not present a Cdh3-based deep cell-cell junctions during *Xenopus* gastrulation. Indeed, vinculin is strongly implicated in E-cad mediated CE of the *Drosophila* epithelial cells (Kale et al., 2018). Combined with our recent exploration of Cdh3 clustering during CE (Huebner et al., 2021), our work here suggests that the paradigm of Cdh1/E-cad-based epithelial cell-cell adhesion cannot necessarily be extending to Cdh3-based mesenchymal cell-cell adhesion. It worth noting, therefore that mammalian Cdh3 (aka P-cadherin) is implicated in collective cell movements in the skin and mammary gland (Cetera et al., 2018; Radice et al., 1997), so further exploration of the cell biological mechanisms of action in those contexts will be exciting.

## Acknowledgements

We would like to acknowledge Dr. Daniel J. Dickinson for critical reading of this manuscript. Work in the JBW lab was supported by grants from the NICHD (RO1 HD099191; R21 HD103882). E.M.M. additionally support from the Welch Foundation (F1515) and the NIGMS (R35 GM122480).

## Supplemental figure legends

**Supplemental figure 1:**
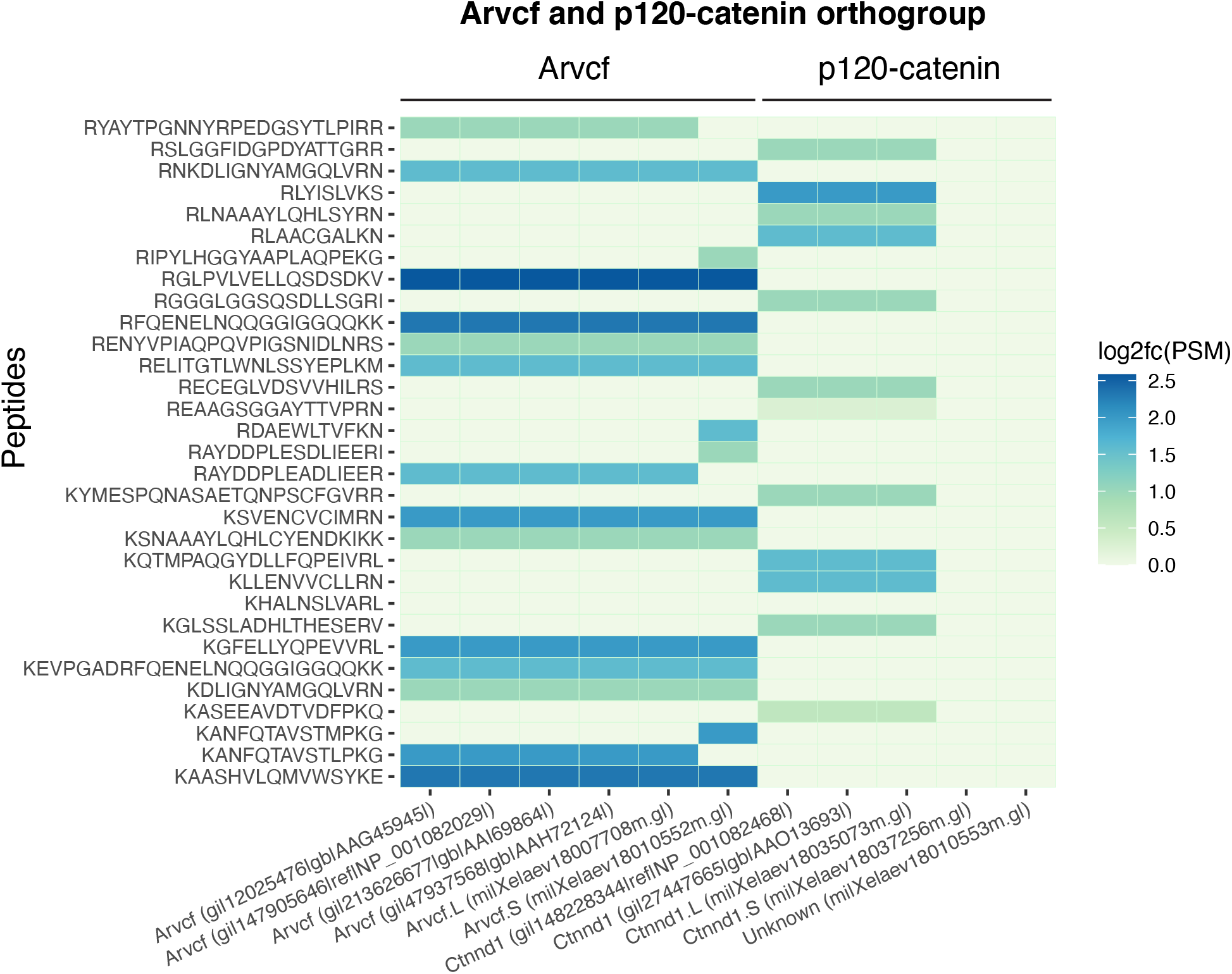
Individual peptide log2fc(PSM) for the ARVCF/p120-catenin orthogroup. **A**. Heatmap showing the peptides identified for the ARVCF/p120-catenin orthogroup. Peptides are on the y-axis and the associated protein is on the x-axis. Color represents the Log2fc(PSM) change comparing cdh3-GFP to the GFP-alone control.

**Supplemental figure 2:**
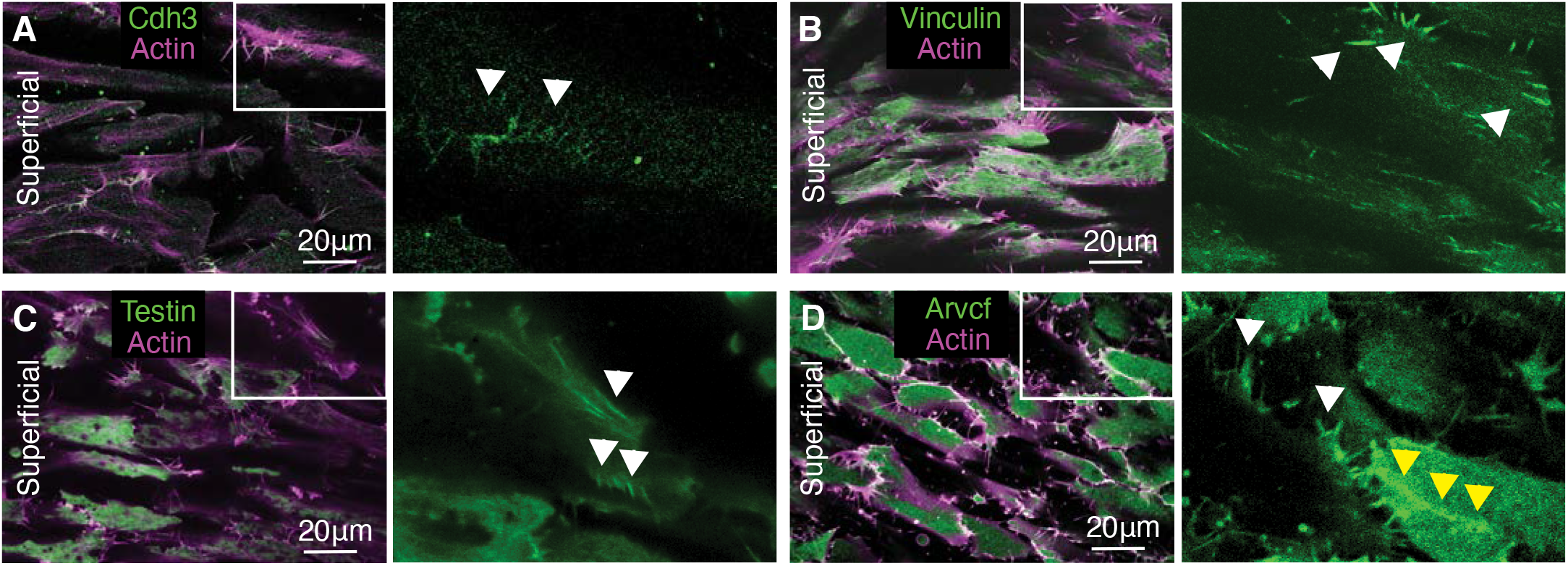
Superficial localization of Cdh3, vinculin, testin, and ARVCF. **A.** Image showing the superficial localization of cdh3-GFP (green) and actin (magenta). Cdh3 can be observed as plaques associated with protrusive structures, white arrowheads. **B.** Image of the superficial localization of vinculin (green) which can be observed in protrusions and in plaque type structures, white arrowheads. **C.** Image of the superficial localization of testin (green). Testin is present in protrusions and superficial plaque structures, white arrowheads. **D.** Image of the superficial localization of ARVCF (green). ARVCF is observed in protrusions, white arrowheads, and is enriched at cell-cell contacts, yellow arrowheads.

**Supplemental figure 3:**
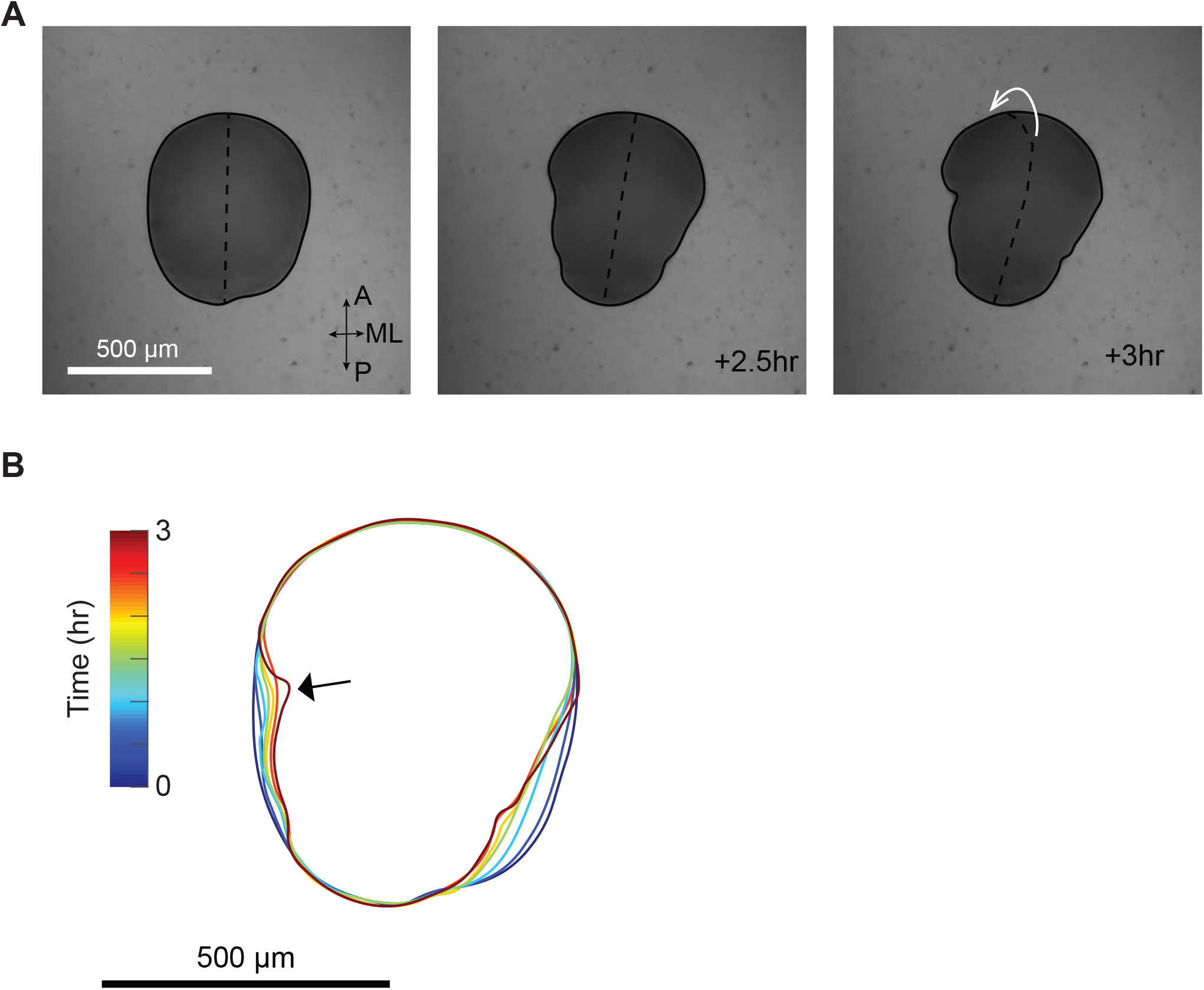
Explant buckling in the semi-rigid gel. **A.** Explant embedded in a semi-rigid gel converged mediolaterally in the first 2.5 hrs and buckled at 3 hr. Black outline, explant-gel interface. Dashed line, explant midline. White arrow, the direction of the out-of-plane buckling. **B.** Outline of the explant during convergent extension. Black arrow, sudden change of the explant shape indicating a buckling happened.

**Supplemental figure 4:**
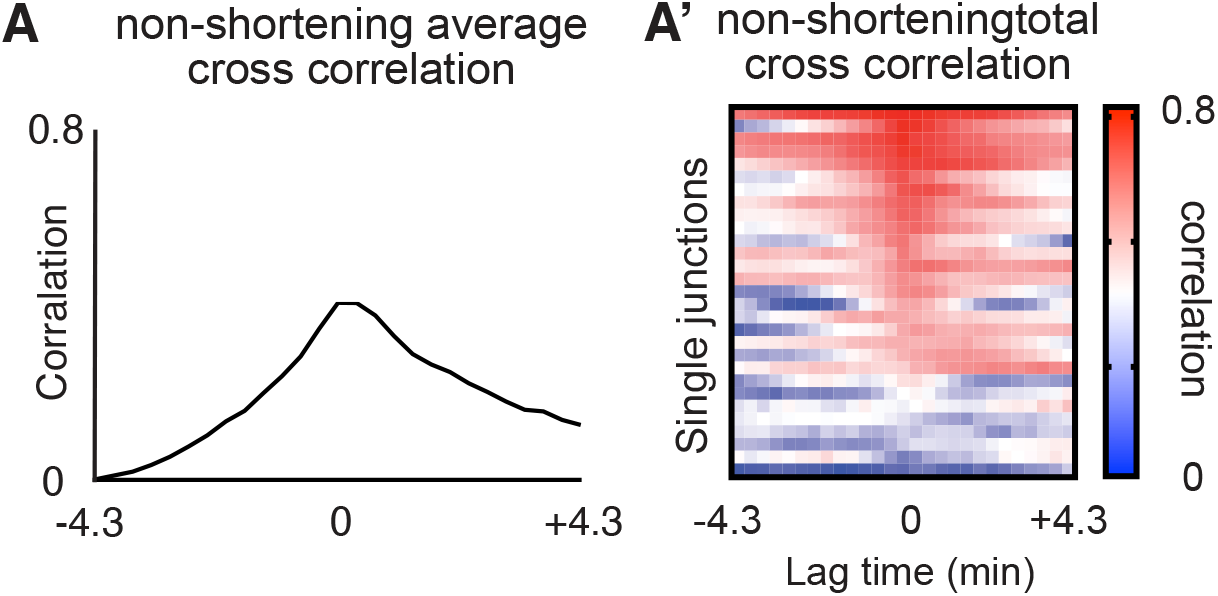
**A**. Graph showing the mean cross correlation between Cdh3 and actin at wildtype non-shortening junctions. **A’**. Heatmap displaying the cross correlations for each individual wildtype non-shortening junctions.

## Methods

### *Xenopus* embryo manipulations

Adult *Xenopus* were maintained in a recirculating aquatic system and female *Xenopus* were ovulated at a maximum of once every three months to ensure high quality embryos. Ovulation is induced by injection of 600 units of human gonadotropin and ovulating females are incubated at 16oC overnight. Following incubation ovulating females produced eggs for ~8 hours. Eggs were acquired by gently “squeezing” ovulating females and then immediately fertilized using crushed testes. Embryos were reared in 1/3X Marc’s Modified Ringer’s (MMR) solution unless otherwise noted. Eggs were dejellied in 3% cysteine (pH 8) for 10 minutes and washed in 1/3X MMR prior to further manipulation.

For microinjection embryos were placed in 2% ficoll in 1/3X MMR 10 minutes prior to injection and then returned to 1/3X MMR 30 minutes after microinjection. Embryos were injected using a Parker’s Picospritizer III with an MK1 manipulator. Embryos were injected in dorsal blastomeres to target the presumptive mesoderm and were injected at the 4- or 32-cell stage as noted in the manuscript.

Microdissections were performed in Steinberg’s solution and explants were maintained in Steinberg’s after dissection. Dissections were performed under a stereoscope using eyelash knives and hair loops. Embryos were dissected at st.~10.25 (Keller explants (Sive et al., 2007)) for mass spec experiments and for imaging experiments. Embryos were dissected at ~st.12 (dorsal explants (Zhou et al., 2015)) for biomechanical assays and the later stage explants were used here because earlier stage explants were not sufficiently stiff for biomechanical manipulations and measurements.

### Morpholino, antibodies, plasmids, and cloning

The Arvcf morpholino was previously characterized (Cho et al., 2011; Fang et al., 2004) and was ordered from Gene Tools. The Arvcf polyclonal antibody was also previously developed and characterized (Paulson et al., 2000) and was gifted from the laboratory of Pierre D. McCrea. Cdh3 antibody was ordered from the Developmental Studies Hybridoma Bank (catalog number 6B6). The α-catenin antibody was purchased from Sigma (catalog number C2081). Cdh3-GFP (Pfister et al., 2016), lifeact-RFP, membrane-GFP, and membrane-BFP were made in pCS105. Vinculin-GFP was made in pCS108 and was a gift from the Paris Skourides lab. Myl9-GFP was made in pCS107 (Shindo et al., 2019). The *Xenopus* Arvcf and testin sequences were obtained from www.xenbase.org and the open reading frames were amplified from a *Xenopus* cDNA library. The Arvcf and testin ORFs were then inserted into the pCS10R-GFP vector.

Arvcf MO: 5’-ACACTGGCAGACCTGAGCCTATGGC -3’

### Morpholino and mRNA microinjections

Capped mRNA was produced using Thermo Fisher SP6 mMessage mMachine kit (catalog number AM1340). mRNAs were injected at the following concentrations vinculin-GFP (50pg), testin-GFP (50pg), Arvcf for imaging and rescue (50pg), lifeact-RFP (100pg), membrane-GFP for imaging and mass-spec (100pg), membrane-BFP (100pg), Myl9-GFP (50pg), Cdh3-GFP for imaging and mass-spec (50pg). Arvcf morpholino was injected at a concentration of 30ng.

### Immunoprecipitation of *Xenopus* Keller explants for mass-spectrometry

Embryos were injected in the presumptive dorsal mesoderm at the 4-cell stage with either Cdh3-GFP or GFP alone. Keller explants were then excised from st.10.25 embryos and explants elongated to ~st.12. Explant immunoprecipitation (IP) was then performed using a GFP-Trap Agarose Kit (ChromoTek, catalog number gtak-20) and proteins were eluted in 2X sample buffer. The IP experiment was performed in 2 replicates with approximately 800 explants per condition per replicate.

### Affinity purification-mass spectrometry

Immunoprecipitated proteins were prepared for mass spectrometry as described in (Lee et al., 2020). Mass spectrometry was performed on a Thermo Orbitrap Fusion for the first and second injections of the first biological replicate of Cdh3-GFP and on a Themo Orbitrap Fusion Lumos Tribrid for the third and fourth injections. Both injections of the second biological replicate of Cdh3-GFP were performed on a Thermo Orbitrap Fusion. In all cases, peptides were separated using revere phase chromatography on a Dionex Ultimate 3000 RSLCnano UHPLC system (Thermo Scientific) with a C18 trap to Acclaim C18 PepMap RSLC column (Dionex; Thermo Scientific) configuration and eluted using a 3% to 45% gradient over 60 min. with direct injection into the mass spectrometer using nano-electrospray. For the second biological replicate and the second two injections of the first biological replicate, data were collected using a data-dependent high energy-induced dissociation (HCD) method. The first and second injections of the first biological replicate were collected using a collision-induced dissociation (CID) method. Full precursor ion scans (MS1) were collected at 120,000 m/z resolution, and monoisotopic precursor selection and charge-state screening were enabled using Advanced Peak Determination (APD), with ions of charge ≥ +2 selected for HCD with stepped collision energy of 30% +/− 3% (Lumos) or 31% +/− 4% (Orbitrap Fusion) or 35% collision energy for CID.

### Protein interaction analysis

Raw MS/MS spectra were processed using Proteome Discoverer (v2.3). We used the Percolator node in Proteome Discoverer to assign unique peptide spectral matches (PSMs) at FDR < 5% to the composite form of the *X. laevis* reference proteome described in (Drew et al., 2020) which comprises both genome-derived Xenbase JGI v9.1 + GenBank *X. laevis* protein sets, but with homeologs and highly related entries combined into eggNOG vertebrate-level orthology groups (Huerta-Cepas et al., 2017), based on the method developed in (McWhite et al., 2020). To identify proteins statistically significantly associated with Cdh3, we calculated both a log2 fold-change and a Z-score for each prey protein based on the observed PSMs in the bait versus control pulldown. Calculations for the fold-change and Z-score were performed as described in (Lee et al., 2020) and (Lu et al., 2007). We determined significance by calculating p-values for each Z-score using the pnorm distribution function available in the R Stats Package (v3.6.1). We corrected for multiple comparisons by computing the Benjamini-Hochberg false discovery rate using the p.adjust function, also from the R Stats Package (v3.6.1). Probability values and false discovery rates are provided in data table 1.

### Mass spectrometry data deposition

Proteomics data has been deposited into MassIVE which in turn was passed to ProteomeXchange. The MassIVE accession # is MSV000087312 (data available for download at https://ftp://massive.ucsd.edu/MSV000087312/). The ProteomeXchange # is PXD025665 (http://proteomecentral.proteomexchange.org/cgi/GetDataset?ID=PXD025665-1).

### Cellular scale imaging of Xenopus mesodermal cells

For a detailed description of high magnification Xenopus imaging please see (Kieserman et al., 2010). Briefly, Keller explants were mounted on fibronectin coated coverslips, mesoderm facing the coverslip, and held in place with a glass “chip” and vacuum grease. Explants were incubated at room temperature for 4 hours or at 16°C overnight. Confocal images were acquired using either a Nikon A1R or a Zeiss LSM700 (40x or 60x lens). Time-lapse movies were acquired with a 20 second time interval. Images were either acquired at the superficial surface (at the coverslip/cell interface) or at the deep cell surface (5μm above the coverslip).

### Measurement of protein intensities at deep cell junctions (intensity plots)

Image analysis was performed using the open-source image analysis software Fiji (Schindelin et al., 2012). Images were first processed using a 50-pixel rolling ball radius background subtraction. Then a 10μm straight line was drawn across each cell-cell junction with the line centered on the junction. The line was then converted to a region of interest and the Multi Plot tool was used to extract intensity values across the region of interest. Intensity plots were then statistically compared using a Kolmogorov-Smirnov test.

### Imaging and measurement of embryos and dorsal explants

Whole embryos and dorsal explants images were acquired using a Zeiss AXIO Zoom stereoscope. Embryos, for the knockdown and rescue experiment, were kept at room temperature until stage 40 and then fixed with MEMFA in glass vials for 1 hour. Post fixation samples were washed three times in 1X PBS and images were acquired. The embryo anterior-posterior length was then measured using the line and measurement tools in Fiji and statistically compared using an ANOVA test. Dorsal explants were dissected from late gastrula embryos (~st.12), allowed to heal for 30 minutes, and pre-CE images were acquired. Explants were then incubated for 4 hours at room temperature and post-elongation images were acquired. Explant lengths were measured using Fiji and statistically compared using a Student’s t-test.

### Immunostaining Keller explants and quantification of endogenous Arvcf and Cdh3 in knockdown cells

Embryos were injected at the 4-cells stage in a single dorsal blastomere with Arvcf morpholino and membrane BFP to generate animals with a mosaic Arvcf knockdown. Keller explants were then dissected from early gastrula embryos (~st.10.25) and mounted on fibronectin coated coverslips. Samples were incubated at room temperature for four hours or overnight at 16°C. Explants were then fixed in 4% paraformaldehyde for 1 hour and washed 3 times in PBS to remove fixative. Next samples were permeabilized with 0.05% Triton X-100 in PBS for 30 minutes and blocked in 1% Normal Goat Serum (NGS) in PBS for 2 hours at room temperature. The primary antibody, for both Arvcf and Cdh3, was diluted 1:100 in 1% NGS/PBS and explants were incubated in primary antibody overnight at 4°C. Samples were then blocked a second time at room temperature for 1 hour and then washed twice with fresh blocking solution. Secondary antibody (goat anti-Mouse 488, #A32723) was diluted 1:500 and samples were incubated at 4°C overnight. Finally, samples were washed three times in 1X PBS and imaged. Images were processed using a 50-pixel rolling ball radius background subtraction. Then the Fiji segmented line tool was used to create a region of interest circling the plasma membrane of wildtype or Arvcf knockdown cells. The protein mean intensity was then measured and samples were statistically compared using a Student’s t-test.

### Constrained explant elongation assay

We used semi-compliant agarose gel to provide external mechanical constraint to CE. We used 0.3% low-melting-point (LMP) agarose (Promega, Madison, WI) gel freshly made in the Steinberg’s solution and kept at 37 °C to remain liquid. Explants were dissected from late gastrula embryos (~st.12) and allowed to heal and clear debris for 30 min. Explants were then rinsed twice with liquid gel cooled down to RT and transferred to individual culture wells casted with 2% agarose gel and pre-filled with liquid LMP agarose gel. Once explants were positioned and oriented, culture wells were moved to a 13.5 °C incubator for 20 min to solidify the LMP agarose gel, then moved back to RT. Explants were then incubated for 4 hours to allow CE.

### Tissue scale force measurement and tissue stiffness estimation

We adapted a previously reported method for the biomechanical testing of the explants during CE (Zhou et al., 2015). We embedded explants dissected from late gastrula embryos (~st.12) in 0.6% LMP agarose gel instead. 0.6% LMP agarose gel was the softest one that limited the induced maximum strain below 10% within 3-hour incubation, which enabled a linear viscoelastic finite element model for force estimation. 0.5 μm red fluorescent latex beads (Sigma) were evenly dispersed in the liquid LMP agarose gel as markers for tracking gel deformation. After gel solidified, the fluorescent beads were scanned during CE with a confocal microscope (Nikon A1R or Zeiss LSM 700, 10x objective). Z-stacks were collected over 40 μm near the dorsal-ventral mid-plane every 20 min.

We observed both wildtype and KD explants embedded in 0.6% agarose gel buckled at the end of the 3-hour incubation. A sudden bend of an explant at the AP poles (“buckling”) was also associated with reduced gel deformation and redirected explant elongation. We took the last frame before explant buckling to quantify the maximum explant elongation force. We first created maximum z-projections of the beads image and aligned it to the one from the first time point using ImageJ (NIH) and analyzed bead displacement using PIVlab (Thielicke, Retrieved February 14, 2020; Thielicke and Stamhuis, 2014; Zhou et al., 2015). We then applied finite element analysis (FEA) to estimate the explant elongation force using customized MATLAB scripts. We assumed that all forces on the gel were applied at the gel-tissue interface and were compressive. The gel was modeled as an isotropic, linear viscoelastic material with bulk elastic modulus of 30 Pa and Poisson’s ratio of 0.5 (Normand et al., 2000; Zhou et al., 2015). Forces were estimated by iterations that minimized the difference between a simulated displacement field and the measured displacement filed from bead tracking. The Von Mises stresses were then calculated for display. The maximum compressive (2^nd^ principle) stresses and average stress along the AP axis were calculated for statistics.

We next used the maximum elongation force, which was also the critical buckling force to estimate tissue stiffness. We simplified explant as an isotropic, linear viscoelastic column with a rectangular cross section and assumed the reactive tissue extending force applied a uniform load along the AP axis. Euler’s buckling load give the formula *F_cr_* = *π*^2^*EI*/*L*^2^, where 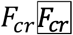 is Euler’s critical load, E is the Young’s modulus of the explant, *I* is the minimum area moment of inertia of the cross section, and *L* is the length of the explant in the AP axis. We can also express *I* as *I* = *t*^3^*w* / 12, where 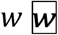 is the width of the explant, and *t* is the thickness. With the assumption of uniform loading, *F_cr_* = *F_s_t* / *t_s_*, where *t_s_* is the scanned thickness and *F_s_* is the estimated force from FEA. Thus, the tissue stiffness can be expressed as *E* = 12*F_s_L*^2^ / *π*^2^*wt*^2^*t_s_*.

### Measurement of cell polarization

Embryos were injected and Keller explants were prepared as described above. We then used the Fiji straight line tool to measure the mediolateral length and anterior-posterior width of individual cells. We then report the cell polarization as the ratio of the mediolateral length over the anterior-posterior width. Samples were statistically compared using a Student’s t-test.

### Measurement of Cdh3 and actin oscillations

Images were processed with 50-pixel rolling ball radius background subtraction. The Fiji segmented line tool, with width set to the thickness of the junction (~16 pixels), was used to set a line of interest (LOI) across the length of the cell junction. Next the multi-plot tool was used to extract cdh3 intensity values across the length of the cell junction and the measure tool was used to collect data such as junction length and mean intensity values. The Fiji Time Lapse plugin Line Interpolator Tool was used to make successive measurements for movies. Here a segmented line LOI was drawn every 10-30 frames, the line interpolator tool was then used to fill in the LOIs between the manually drawn LOIs allowing rapid semi-manual segmentation. The multi-plot tool and measure tool were then used to extract data for each time-point of the movie.

## Notes

### Competing Interest Statement

The authors have declared no competing interest.

